# Size, not temperature, drives cyclopoid copepod predation of invasive mosquito larvae

**DOI:** 10.1101/839480

**Authors:** Marie C. Russell, Alima Qureshi, Christopher G. Wilson, Lauren J. Cator

## Abstract

During range expansion, invasive species can experience new thermal regimes. Differences between the thermal performance of local and invasive species can alter species interactions, including predator-prey interactions. The Asian tiger mosquito, *Aedes albopictus*, is a known vector of several viral diseases of public health importance. It has successfully invaded many regions across the globe and currently threatens to invade regions of the UK where conditions would support seasonal activity. We assessed the functional response and predation efficiency (percentage of prey consumed) of the cyclopoid copepods *Macrocyclops albidus* and *Megacyclops viridis* from South East England, UK against newly-hatched French *Ae. albopictus* larvae across a relevant temperature range (15, 20, and 25°C). Predator-absent controls were included in all experiments to account for background prey mortality. We found that both *M. albidus* and *M. viridis* display type II functional response curves, and that both would therefore be suitable biocontrol agents in the event of an *Ae. albopictus* invasion in the UK. No significant effect of temperature on the predation interaction was detected by either type of analysis. However, the predation efficiency analysis did show differences due to predator species. The results suggest that *M. viridis* would be a superior predator against invasive *Ae. albopictus* larvae due to the larger size of this copepod species, relative to *M. albidus*. Our work highlights the importance of size relationships in predicting interactions between invading prey and local predators.

## Introduction

An invasion of the Asian tiger mosquito, *Aedes albopictus*, into the UK is a major public health concern. This species is not only an aggressive nuisance biter, but also a known vector of arboviruses such as dengue, chikungunya, yellow fever, and Zika (Centers for Disease and Prevention, 2017). The ability of *Ae. albopictus* mosquitoes to lay desiccation-resistant eggs has enabled their introduction into cities across the globe, often via used tire shipments (Eritja et al., 2005, Juliano and Lounibos, 2005, Lounibos, 2002, Medlock et al., 2006, Benedict et al., 2007). According to the European Centre for Disease Prevention and Control, *Ae. albopictus* populations have already established throughout the vast majority of Italy, southern France, and as far north as the French department of Aisne (European Centre for Disease and Control, 2019). Although this species has not yet established in the UK, it has been introduced in Kent, a coastal county in the southeast of the UK, where 37 *Ae. albopictus* eggs were found in September of 2016 (European Centre for Disease and Control, 2019, Medlock et al., 2017).

Approximately one decade prior to the first recorded introduction of *Ae. albopictus* into the UK, a model based on abiotic factors including photoperiod, rainfall, and temperature suggested that upon introduction, *Ae. albopictus* adults would be active from May to September in areas near London and the southern coastal ports (Medlock et al., 2006). A more recent study that also considered diurnal temperature range and human population density found that the most suitable areas for *Ae. albopictus* in the UK currently are centered around London (Metelmann et al., 2019). Owing to the warming climate, the UK is expected to become increasingly suitable for *Ae. albopictus* establishment; projections of climate and human population conditions into future decades suggest that if introduced, the vector species could establish throughout most of England and southern Wales during the 2060s (Metelmann et al., 2019). Another recent model predicts that the UK will report *Ae. albopictus* presence by either 2050 or 2080, depending on the patterns of future greenhouse gas emissions (Kraemer et al., 2019).

Predation by copepods has been identified as a biological method for controlling *Ae. albopictus* in Europe following the success of past field trials (Baldacchino et al., 2015). The cyclopoid copepod *Macrocyclops albidus* has previously been used to eliminate *Ae. albopictus* populations from tire piles in New Orleans, Louisiana, USA (Marten, 1990a). In 1994, *Mesocyclops aspericornis* were distributed into wells in Charters Towers, Queensland, Australia to eliminate *Ae. aegypti* populations (Russell et al., 1996). In addition, *Mesocyclops* copepods were used to effectively control populations of *Ae. aegypti* and *Ae. albopictus* in six communes in northern Vietnam (Kay et al., 2002). A semi-field study conducted in Bologna, Italy in 2007 found that *M. albidus* reduced the density of *Ae. albopictus* in experimental drums by greater than 99% (Veronesi et al., 2015). In the UK, laboratory experiments using cyclopoid copepods from Northern Ireland against *Culex pipiens* mosquito larvae from Surrey, UK and *Ae. albopictus* larvae from Montpellier, France have supported the use of copepods as biocontrol agents (Cuthbert et al., 2019b, Cuthbert et al., 2018b). Previous work indicates that the attack rates of *M. albidus* and *Megacyclops viridis* predators against *Cx. pipiens* prey tend to increase with temperature (Cuthbert et al., 2018b). However, the use of UK copepods as predators of invasive *Ae. albopictus* larvae has not been thoroughly examined over the range of temperatures that the invasive larvae are predicted to experience. Due to the potentially negative impacts of exporting copepods to non-native regions for biocontrol purposes (Coelho and Henry, 2017), it is important to investigate the performance of copepods local to the predicted sites of *Ae. albopictus* establishment: London and South East England.

Populations of *M. albidus* and *M. viridis* cyclopoid copepods from the benthos of the Cumbrian lakes in the UK have previously been studied at temperatures from 5 to 20°C (Laybourn-Parry et al., 1988). Based on the dry body masses of adult specimens, males were consistently smaller than females, and *M. albidus* copepods were consistently smaller than *M. viridis* (Laybourn-Parry et al., 1988). The general adult body length ranges for *M. albidus* and *M. viridis* are 1.3 – 2.5 mm (Einsle, 1993) and 1.2 – 3 mm (Dussart, 1969, Einsle, 1988, Kiefer, 1960), respectively. These are small enough to enable the distribution of copepod cultures to mosquito larval habitats using “a simple backpack sprayer with a 5 mm hole in the nozzle,” as has been previously recommended (Marten, 1990b). Copepods reproduce sexually, and females that can produce new egg sacs every 3-6 days tend to predominate in mature populations (Marten and Reid, 2007). Cyclopoid copepods are considered sit-and-wait ambush predators because of their attack behaviors, which have been described in six steps: encounter, aiming, stalking, attack, capture, and ingestion (Awasthi et al., 2012). When preying on mosquito larvae, copepods often avoid ingesting the head and thorax (Awasthi et al., 2012).

Functional response curves were originally developed to relate the number of prey attacked by an invertebrate predator to the prey density (Holling, 1959, Holling, 1966). Previous work has suggested that the best predators to use in an “inundative release” biocontrol program are those that have type II functional responses to prey density, so that the predators’ attack rates are high even at low prey densities (Daane et al., 1996). Predators that display a type III functional response against an invasive prey species —such as signal crayfish (*Pacifastacus leniusculus*) against New Zealand mud snails (*Potamopyrgus antipodarum*)—may be able to limit the spread of the invader, but cannot prevent it from establishing (Twardochleb et al., 2012). Cyclopoid copepods from the UK have previously exhibited type II functional response curves when provided with *Ae. albopictus* prey at a single temperature setting (Cuthbert et al., 2019b).

A meta-analysis of fifty functional response curves of cyclopoid copepod predators showed a “monotonically increasing effect of temperature” on the attack rate parameter (Kalinoski and DeLong, 2016). In addition, gut clearance rate, another measure of foraging rate, has been demonstrated to increase linearly with temperature in planktonic copepods (Dam and Peterson, 1988, Irigoien, 1998). However, five data points that were excluded from the linear regression analysis and described as “distinctly separate” would have suggested a decrease in gut clearance rate at higher temperatures (Irigoien, 1998). Furthermore, recent literature shows that the temperature dependence of attack rate is unimodal, or “hump-shaped,” especially when the temperature range exceeds thermal optima, and suggests that the impact of climate change on predation will depend on the thermal optima of consumers (Englund et al., 2011, Uiterwaal and Delong, 2020).

If UK cyclopoid copepods are used as biocontrol agents against *Ae. albopictus*, they are likely to experience temperatures higher than those of their natural habitats not only because of climate change, but also because the predation interaction would occur mostly in artificial containers, where low volumes of water can warm more quickly, relative to the warming expected in lakes and ponds. The usual environmental temperatures experienced by *M. albidus* and *M. viridis*, 5 to 20°C (Laybourn-Parry et al., 1988), are expected to influence their thermal optima (Dell et al., 2011). An analysis of 2,083 functional responses demonstrated that the thermal optima of attack rate and handling time parameters often fall within the range of 15-25°C (Uiterwaal and Delong, 2020). Among 69 unimodal thermal responses of freshwater organisms, such as cyclopoid copepods, a mean optimal temperature of 21°C was observed (Dell et al., 2011). However, the thermal optimum of the prey species can also influence the temperature dependence of predation interactions. The optimal temperature for French *Ae. albopictus* is likely to be higher than that of UK copepods and has been recorded at 29.7°C for immature *Ae. albopictus* originating from La Réunion, an island in the southwest region of the Indian Ocean (Delatte et al., 2009). Such asymmetries in the thermal response of consumer and prey species may modify the effect of temperature on the predation interaction (Dell et al., 2014, Grigaltchik et al., 2012).

In this study, we determined the type of functional response of *M. albidus* and *M. viridis* copepods from Surrey, UK against *Ae. albopictus* larvae from Montpellier, France at three different temperatures (15, 20, and 25°C) likely to be experienced by container-dwelling mosquito larvae in London and South East England. We also calculated predation efficiency, the percentage of larvae consumed, of both copepod species at a constant mosquito prey density under the same three temperatures.

## Materials and methods

### Collection of field temperature data

Previous work has suggested that the microclimates experienced by *Ae. albopictus* are not represented well by climate data reported from local weather stations (Murdock et al., 2017). Since used tire shipments are a common source of *Ae. albopictus* introductions (Eritja et al., 2005, Juliano and Lounibos, 2005, Lounibos, 2002, Medlock et al., 2006, Benedict et al., 2007), we measured the temperature of rainwater collected in used car tires that were divided between an urban London site and a suburban site. These temperature data provided rough guidelines for our choice of the three temperatures to be tested in our functional response and predation efficiency experiments. The minimum water temperature was 9°C, the 25^th^ percentile was 16.1°C, the median was 18.1°C, the 75^th^ percentile was 20°C, and the maximum was 25°C. Thus, our choice of 15, 20, and 25°C as the three main temperatures of interest is representative of more than 75% of the tire water temperatures recorded in the field from May through September of 2018 (Supporting Information).

### Local copepod cultures

Adult gravid female copepods were collected in August of 2018 from the edge of Longside Lake in Egham, Surrey, UK (N 51° 24.298’, W 0° 32.599’) using a 53 μm sieve (Reefphyto Ltd, UK). Separate cultures were started from each gravid female. The copepods were kept in 3 L containers of spring water (Highland Spring, UK) at a 12:12 light/dark cycle, and 20 ± 1°C, a temperature previously found to increase the portion of their life cycle spent in the reproductive phase (Laybourn-Parry et al., 1988). *Chilomonas paramecium* and *Paramecium caudatum* (Sciento, UK) were provided *ad libitum* as food for the copepods (Suarez et al., 1992). The ciliates were cultured in 2 L flasks containing boiled wheat seeds; boiled wheat seeds were also added to the copepod containers (Suarez et al., 1992). Adult copepods were identified as *Macrocyclops albidus* (Jurine, 1820) and *Megacyclops viridis* (Jurine, 1820) by Dr. Maria Hołyńska from the Museum and Institute of Zoology in Warsaw, Poland.

### Temperate *Ae. albopictus* colony care

A colony of *Ae. albopictus* mosquitoes (original collection Montpellier, France 2016 obtained through Infravec2) was maintained at 27 ± 1°C, 70% relative humidity, and a 12:12 light/dark cycle. The colony maintenance temperature of 27°C was chosen because it falls at the upper bound of the 95% confidence interval that was fitted around mean daily Montpellier temperatures from 1996-2005 in the summer months, and these temperatures are expected to increase in the coming decades (Zidon et al., 2016). Larvae were fed fish food (Cichlid Gold Hikari^®^, Japan), and adults were given 10% sucrose solution and horse blood (First Link Ltd, UK) administered through a membrane feeding system (Hemotek^®^, Blackburn, UK).

### Design of functional response experiments

Adult non-gravid female copepods, identified by larger relative size, were removed from their culture and each was placed in a Petri dish (diameter: 50 mm, height: 20.3 mm) holding 20 mL of spring water. At approximately 11am, the copepods were placed in three different controlled environments set to 15 ± 1, 20 ± 1, and 25 ± 1°C, all at a 12:12 light/dark cycle, to begin a 24 h starvation and temperature-acclimation period for the predators (Fig. 1). Each combination of copepod species (*M. albidus* or *M. viridis*) and temperature had 28 copepods, each one held in its own Petri dish; there were 168 copepods in total (Fig. S3a). Ten additional gravid females were randomly selected from each species of copepod and preserved in 80% ethanol for size measurements.

**Fig. 1.**
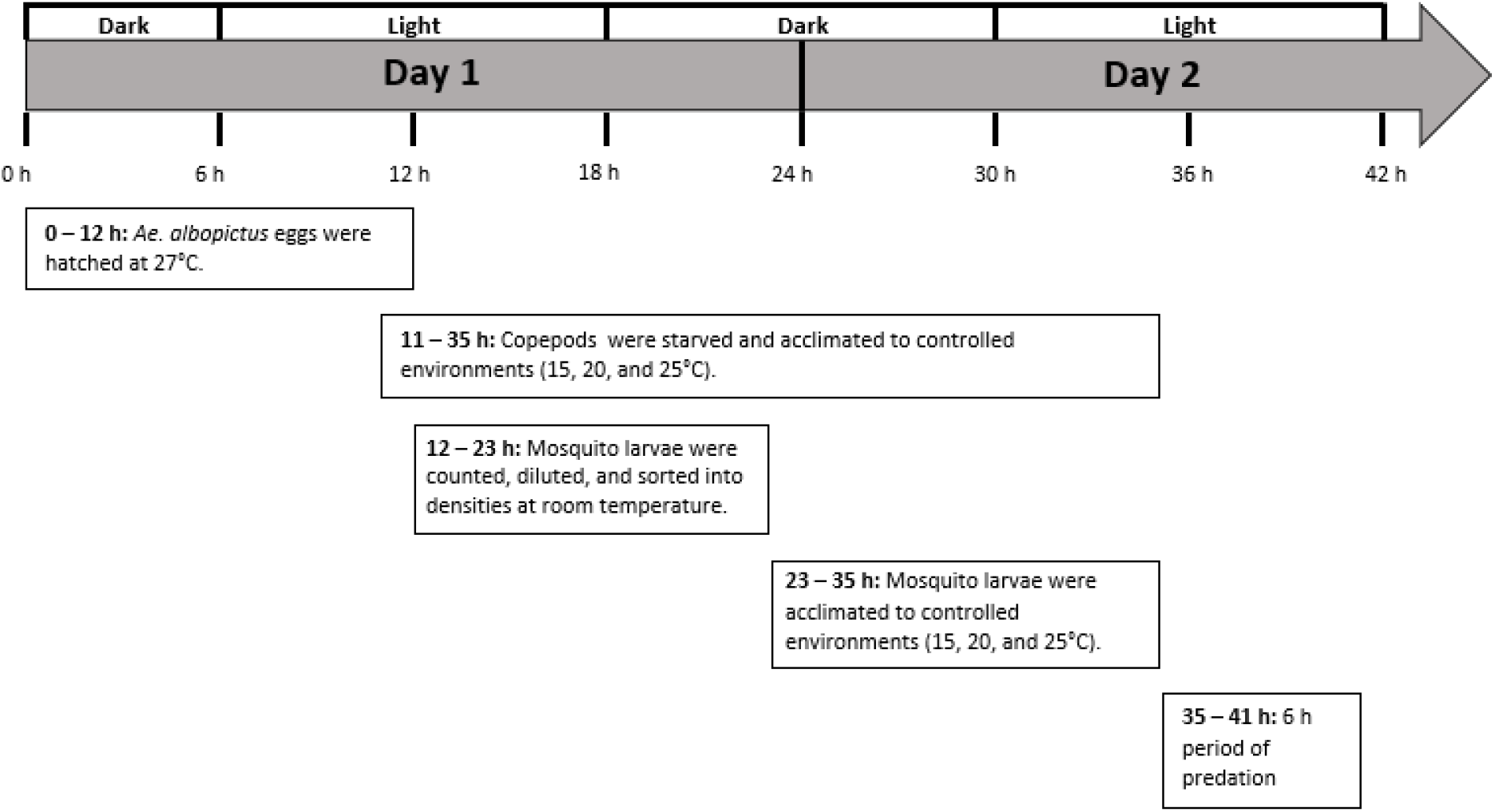
General schedule of both functional response and predation efficiency experiments

At noon, groups of 87 newly-hatched *Ae. albopictus* larvae (Supporting Information) were each pipetted into a small plastic tub containing 150 mL of spring water to strongly dilute any residual food from the hatching media. Each tub of 87 larvae was then split into seven Petri dishes each containing 20 mL of spring water and the following larval densities: 1, 2, 4, 8, 16, 24, and 32. All larvae were counted into dishes at room temperature and then divided into the three different controlled environments, allowing a 12 h temperature acclimation period before predator introduction (Fig. 1). For each combination of copepod species and temperature, there were 35 Petri dishes containing a total of 435 newly-hatched *Ae. albopictus* larvae, with a total of 2,610 larvae used across all predator treatments and controls (Fig. S3a). This sample size allowed for four replicates of each larval density and one predator-absent control at each density, for each combination of copepod species and temperature (Fig. S3a).

At approximately 11am the next day, the copepod predators held at three different temperatures were introduced to larval Petri dishes of matching controlled temperatures (Fig. 1). The copepods were removed after a 6 h period of predation (Fig. 1), which follows the schedule of similar experiments (Cuthbert et al., 2018a, Cuthbert et al., 2019a). Each copepod was preserved in 80% ethanol so that its body size could later be measured and matched to the larval count data. Immediately following the removal of the copepods, the number of surviving larvae in each Petri dish was counted and recorded. In addition, the body lengths of the 10 gravid females selected from each species of copepod, as well as the body lengths of all copepods included as predators in the functional response experiments, were measured from the front of the cephalosome to the end of the last urosomite (Dussart and Defaye, 2001).

### Design of predation efficiency experiment

In addition to functional response, predation efficiency has previously been measured to assess copepod predators of mosquito larvae (Baldacchino et al., 2017). We measured predation efficiency using a constant density of 24 larvae in 20 mL of spring water. This density was chosen from the higher end of the range of densities used in the functional response experiments, where the curves generally start to plateau (Juliano, 2001) (Fig. 2a).

**Fig. 2.**
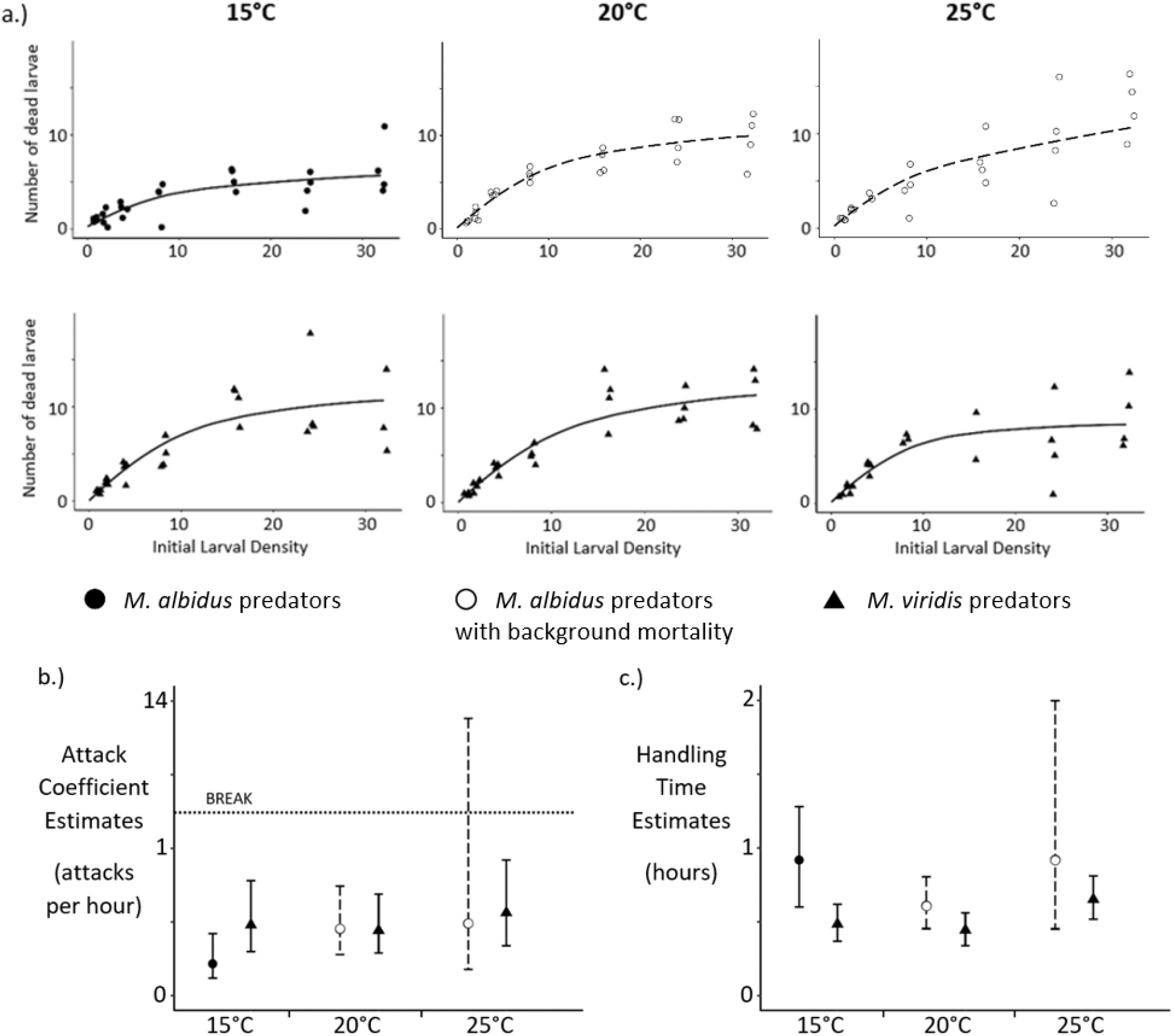
**a.)** Type II functional response curves for each combination of copepod species and temperature (Points are shown in the “jitter” position.) **b.)** Attack coefficient estimates and 95% confidence intervals by copepod species and temperature **c.)** Handling time estimates and 95% confidence intervals by copepod species and temperature

*Ae. albopictus* larvae were hatched (Supporting Information) and any residual food was diluted in spring water following the same procedure used for the functional response experiments. Adult non-gravid female copepods were each placed in a Petri dish at approximately 11am for a 24 h period of starvation and acclimation to three different temperature settings: 15 ± 1, 20 ± 1, and 25 ± 1°C, all at a 12:12 light/dark cycle (Fig. 1). The largest non-gravid copepods of each species were selected to minimize the risk of selecting males or immature stages. The Petri dishes containing larvae were split into the three different temperature settings 12 h prior to the introduction of copepod predators, and there was a 6 h period of predation between 11am and 5pm (Fig. 1). At the end of the 6 h period, the copepods were removed, and each was stored in 80% ethanol and labelled according to its larval Petri dish. The number of surviving larvae in each Petri dish was recorded immediately after removing the copepods.

At each of the three temperature settings, there was a total of 24 Petri dishes, each containing 24 larvae (1,728 larvae across all temperatures); the 24 dishes were divided into three groups (n = 8): one with *M. albidus* predation, one with *M. viridis* predation, and one as a control (Fig. S3b). Every predator treatment was matched to a control that had been held at the same temperature (Fig. S3b). Each temperature setting had a total of 192 control larvae. Larval background mortality was 5% at 15°C, 11% at 20°C, and 3% at 25°C. A total of 24 *M. albidus* and 24 *M. viridis* copepods were used in this experiment. The body lengths of these copepods were measured after each was preserved in 80% ethanol.

### Functional response curve analysis

The general shapes of the functional response curves were determined for each combination of copepod species and temperature using predator-present data. When polynomial logistic functions are fitted to describe the relationship between the proportion of larvae killed and the initial larval density, a negative first-order term indicates a type II response, and a positive first-order term indicates a type III response (Juliano, 2001, Pritchard et al., 2017b). The “frair_test” function (“frair” package, R version 3.4.2) was used to determine the shapes of the functional response curves according to the sign and significance of first-order and second-order terms in logistic regressions (Pritchard et al., 2017a, Pritchard et al., 2017b). Three observations were excluded from the *M. albidus* at 25°C group — two due to copepod deaths during the starvation period, and one because two copepod predators were introduced into the same Petri dish. Three observations were excluded from the *M. viridis* at 25°C group, and one was excluded from the *M. viridis* at 15°C group; all four *M. viridis* exclusions were due to insufficient body length of the copepod indicating that these predators could have been either males or immatures, rather than adult non-gravid females.

Mortality of *Ae. albopictus* larvae was observed in controls for five out of the six combinations of copepod species and temperature. *M. viridis* predators at 25°C was the only group without larval mortality in any of the controls. For *M. albidus*, across all control larvae (n=87) at 15°C, three deaths were observed (3% mortality); four deaths were observed at 20°C (5% mortality); and eleven deaths were observed at 25°C (13% mortality). For *M. viridis*, two deaths were observed among control larvae at 15°C (2% mortality), and three deaths were observed at 20°C (3% mortality). Mortality rates of aedine larvae in predator-absent controls have previously been observed to be as high as 20% (Baldacchino et al., 2017). To account for both background mortality and prey depletion caused by predation, the functional response parameters (attack coefficient and handling time) were estimated by fitting the following differential equation:

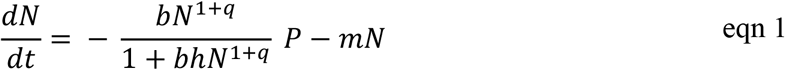

where *dN/dt* is the change in prey density over time, *b* is the attack coefficient, *q* is an exponent that can fall between a type II response (q = 0) and a type III response (q = 1), *h* is the handling time, *P* is the predator density, and *m* is the mortality rate (Rosenbaum and Rall, 2018).

Since background prey mortality should not alter the general shape of the functional response curve (Rosenbaum and Rall, 2018), the *q* parameter was fixed to either 0 or 1 based on the results of the “frair_test” function, which only considers data from predator-present treatments (Pritchard et al., 2017a, Pritchard et al., 2017b). The attack coefficient (*b*) was given a starting value of one, the handling time (*h*) had a starting value equivalent to the inverse of the maximum feeding rate, and the mortality rate (*m*), if applicable, was given a starting value of 0.01 (Rosenbaum and Rall, 2018). The attack coefficient and handling time parameters were estimated based on our empirical data using an iterative maximum likelihood method: “mle2” function, “bbmle” package, R version 3.4.2 (Bolker et al., 2020, Bolker, 2008, Rosenbaum and Rall, 2018).

The “nll.ode.general.mort” function (Rosenbaum and Rall, 2018) was used to fit data that included some background prey mortality to the statistical model (eqn1). In cases where the mortality rate was shown to be insignificant, or where no larval mortality was originally observed, the “nll.bolker” function (Rosenbaum and Rall, 2018), which assumes a mortality rate of zero, was used. Both functions are negative log-likelihood functions that are minimized by applying maximum likelihood estimation methods (Rosenbaum and Rall, 2018). Confidence intervals for the model parameters of interest, attack coefficient and handling time, were computed by the “confint” function, base package, R version 3.4.2. To detect differences in functional response parameter estimates due to either temperature category or copepod species, 95% confidence intervals were compared; this method for comparing parameter estimates across temperatures has previously been used in a similar analysis of copepod predation (Novich et al., 2014).

### Predation efficiency analysis

For each pair of predator treatment and control (Fig. S3b), the copepod’s predation efficiency was calculated according to Abbott’s formula (Abbott, 1987, Baldacchino et al., 2017):

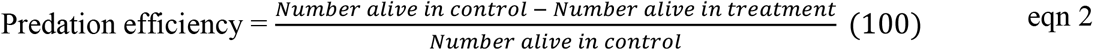

Copepod body lengths, measured in mm, were converted to estimates of body mass (mg) using an equation from previous studies (Alcaraz and Strickler, 1988, Klekowski and Shushkina, 1966, Novich et al., 2014):

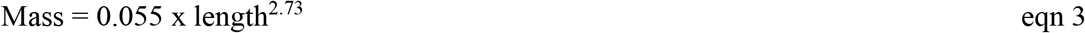

Two linear regression models were fitted to explain predation efficiency. The first tested temperature and copepod species as predictors (eqn S1), and the second tested copepod body mass in place of species (eqn S2). Both models were also fitted without temperature predictors (Supporting Information).

### Body mass difference between copepod species by experimental design

Among the predators used in the functional response experiments, each *M. albidus* copepod was randomly paired with an *M. viridis* copepod (80 pairs). Predation efficiency predators were also randomly paired, yielding 23 pairs of the two species. For each pair, the *M. albidus* body mass was subtracted from that of *M. viridis*, generating 103 differences in body mass due to species. A Wilcoxon rank sum test was used to determine if the body mass differences between copepod species varied by experimental design.

Analyses were completed in R version 3.4.2, and all data will be made accessible from the Dryad Digital Repository.

## Results

### Functional response curves

A type II functional response, demonstrated through logistic regression results (Table S1), was observed in every combination of copepod species and temperature (Fig. 2a). There was a significant effect of background mortality on the change in prey density over time for *M. albidus* at 20 and 25°C (Table S2). Within each copepod species, there was no significant difference in either attack coefficient or handling time due to temperature (Fig. 2b & 2c, Table S4). Within each temperature setting, there was no significant difference in either type of parameter estimate due to copepod species (Fig. 2b & 2c, Table S4). The 95% confidence interval around the attack coefficient estimate for *M. albidus* at 25°C was extremely wide, representing high uncertainty (Fig. 2b). This imprecise estimate was likely due to the highly significant effect of mortality rate (p-value = 0.0040) on the change in prey density over time (Table S2). In addition, the mortality rate’s standard error was higher for *M. albidus* at 25°C than it was for any other data subset that included background larval mortality (Table S2), and the model deviance (−2 log L = 111.25) was higher than that of any other model used to generate parameter estimates (Tables S2 & S3).

### Predation efficiency

Predation efficiency values ranged from 5.0% to 58.3%, with a median of 21.7%, and a slightly right-skewed distribution (skewness = 0.61, kurtosis = 2.9).

A linear model testing copepod species and temperature category as predictors of predation efficiency explained 6.8% of the variance in predation efficiency (Table 1). The predation efficiency of *M. viridis* copepods was 25.7%, significantly higher (p-value = 0.0288) than that of *M. albidus* copepods, which had a predation efficiency of 18.2%, when controlling for differences in temperature (Table 1, Fig. 3a). Interactions between species and temperature were not significant (Species x Temperature_20_, p-value = 0.150; Species x Temperature_25_, p-value = 0.466). Removing the temperature predictors from the model increased the adjusted R^2^ value and decreased the AIC (Tables 3 & S5).

**Table 1.** Linear regression of predation efficiency by copepod species and temperature (n = 47)

| Parameter | Estimate | Standard Error | p-value | Adjusted R<sup>2</sup> | AIC |
| --- | --- | --- | --- | --- | --- |
| Intercept | 18.19 | 3.30 | 1.92 x 10<sup>-6</sup> | 0.068 | 373.1 |
| Species (*M. viridis*, ref = *M. albidus*) | 7.54 | 3.33 | 0.0288 |  |  |
| Temperature (20°C, ref = 15°C) | 0.80 | 4.04 | 0.8432 |  |  |
| Temperature (25°C, ref = 15°C) | 4.56 | 4.10 | 0.2731 |  |  |

**Fig. 3.**
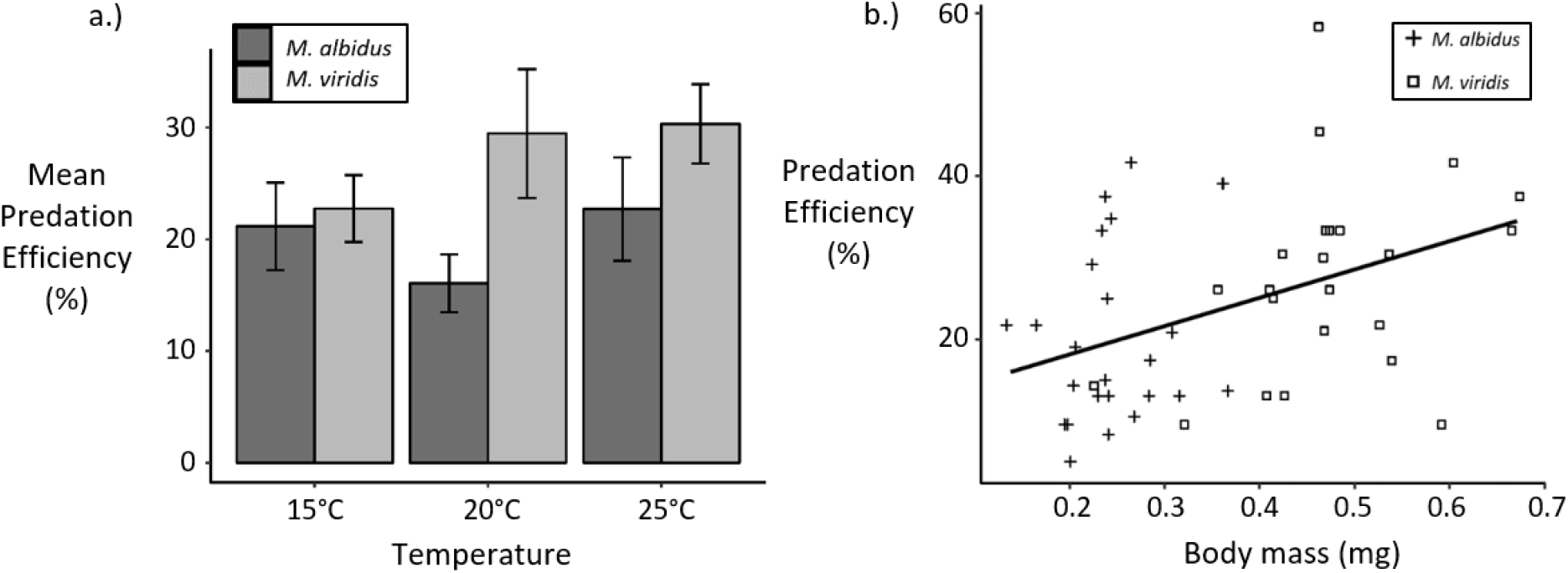
**a.)** Bar chart of predation efficiency by copepod species and temperature (Error bars represent ± the standard error.) **b.)** Predation efficiency by copepod body mass (Points are shown in the “jitter” position.)

A linear model testing copepod body mass and temperature category as predictors of predation efficiency explained 14.0% of the variance in predation efficiency (Table 2). For each 0.1 mg increase in copepod body mass, the predation efficiency increased by approximately 3.4 percentage points (p-value = 0.0042), when controlling for differences in temperature (Table 2, Fig. 3b). Interactions between body mass and temperature were not significant (Mass x Temperature_20_, p-value = 0.611; Mass x Temperature_25_, p-value = 0.565). Removing the temperature predictors from the model improved model fit (Tables 3 & S6).

**Table 2.**
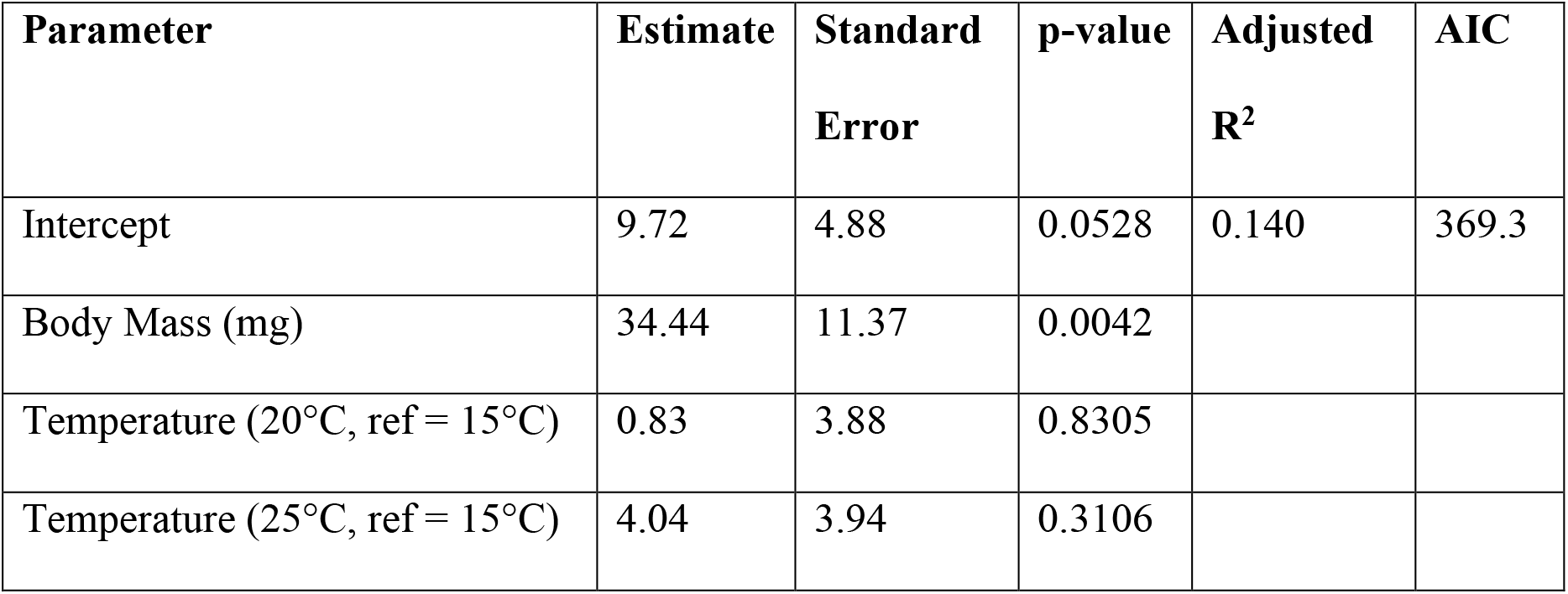
Linear regression of predation efficiency by copepod body mass and temperature (n = 47)

| Parameter | Estimate | Standard Error | p-value | Adjusted R<sup>2</sup> | AIC |
| --- | --- | --- | --- | --- | --- |
| Intercept | 9.72 | 4.88 | 0.0528 | 0.140 | 369.3 |
| Body Mass (mg) | 34.44 | 11.37 | 0.0042 |  |  |
| Temperature (20°C, ref = 15°C) | 0.83 | 3.88 | 0.8305 |  |  |
| Temperature (25°C, ref = 15°C) | 4.04 | 3.94 | 0.3106 |  |  |

**Table 3.** Model selection

| Parameters | Degrees of Freedom | Adjusted R<sup>2</sup> | AIC<sup>a</sup> |
| --- | --- | --- | --- |
| Species, Temperature<sub>20</sub>, Temperature<sub>25</sub> | 43 | 0.068 | 373.1 |
| Species, Temperature<sub>20</sub>, Temperature<sub>25</sub>, Species x Temperature<sub>20</sub>, Species x Temperature<sub>25</sub> | 41 | 0.071 | 390.7 |
| Species | 45 | 0.081 | 362.6 |
| Mass, Temperature<sub>20</sub>, Temperature<sub>25</sub> | 43 | 0.140 | 369.3 |
| Mass, Temperature<sub>20</sub>, Temperature<sub>25</sub>, Mass x Temperature<sub>20</sub>, Mass x Temperature<sub>25</sub> | 41 | 0.107 | 388.8 |
| Mass | 45 | 0.156 | 358.5 |

a.) A difference in AIC value greater than or equal to 2 is the suggested minimum
difference for model selection purposes (Burnham and Anderson, 2002).

### Body mass differences between copepod species by experimental design

*M. viridis* copepods were larger than *M. albidus* copepods in both the functional response and the predation efficiency experiments, and both copepod species were larger in the predation efficiency experiments than in the functional response experiments (Supporting Information, Fig. S4).

The body mass difference between copepod species was greater in the predation efficiency experiments than in the functional response experiments (p-value = 0.0136). In the predation efficiency experiments, the median increase from *M. albidus* to *M. viridis* was 0.24 mg, while in the functional response experiments, the median increase was 0.12 mg.

## Discussion

Both *M. albidus* and *M. viridis* copepods collected from Surrey, UK would be appropriate for use in an inundative release biocontrol program because both predator species display type II functional response curves when presented with invasive *Ae. albopictus* larvae (Table S1). Type II functional response curves are preferred because they demonstrate the predator’s ability to eliminate invasive populations at low prey densities (Daane et al., 1996). Other studies using *M. albidus* and *M. viridis* from Northern Ireland against mosquito larvae prey have also found evidence of type II functional response curves (Cuthbert et al., 2019b, Cuthbert et al., 2018b).

Our functional response analyses did not detect any differences in response type, attack coefficient, or handling time due to copepod species or temperature, tested at 15, 20, and 25°C (Fig. 2, Table S1). In addition, we found that temperature categorical variables were not significant predictors of copepod predation efficiency against *Ae. albopictus* larvae (Tables 1–3). A meta-analysis of 648 functional responses from 86 different studies found that although attack rate increased with temperature, the increase was less steep than that which had been expected based on metabolic theory (Rall et al., 2012). In addition, a previous study of predator-prey interactions involving sit-and-wait copepod predators found no significant difference in the attack parameter of the functional response across three different temperatures: 18, 22, and 26°C (Novich et al., 2014). Since the effect of temperature on sit-and-wait predation is driven mainly by the velocity of the prey (Dell et al., 2014), our results suggest that the velocity of newly-hatched *Ae. albopictus* larvae does not change dramatically over a temperature range of 15-25°C. This is consistent with previous observations of high cold tolerance in aedine larvae. A recent study showed that *Ae. aegypti* larvae from a colony maintained at 22 ± 1°C continued to move in response to probing as the temperature was gradually lowered; the larvae did not stop responding with movement until the temperature reached 6°C, 16°C lower than the temperature at which their colony was maintained (Jass et al., 2019). From a public health perspective, the absence of significantly lower predation efficiency at lower temperatures is useful because *Ae. albopictus* larvae that develop at lower temperatures have been shown to be more susceptible to chikungunya infection as adults (Westbrook et al., 2010).

The term “handling time” has been defined differently by different studies. For example, one study defined it as “the time lapsed between an attack of the prey by the copepod and its resumption of movement after ingestion,” and found that *M. thermocyclopoides* copepods displayed an average handling time of 83 s for first instar *Anopheles stephensi* larvae and 185 s for first instar *Cx. quinquefasciatus* (Kumar and Rao, 2003). These times are consistent with directly-observed ingestion times reported by another study, which states that a mosquito larva is usually consumed by a cyclopoid copepod within a few minutes (Marten and Reid, 2007). However, the original methods designed for fitting functional response curves assume that the predator only has “two time-consuming behaviours – searching and handling of prey” (Holling, 1959). In this context, the “handling time” has been defined as “the time lost from searching per resource consumed,” not as the ingestion time per resource consumed (Uiterwaal and Delong, 2020). Our handling time estimates (Fig. 2c, Table S4) are based on the original framework of functional response analysis, and therefore represent average predator search times per larva consumed. One study found that when directly-observed handling times were adjusted to include both the time that the predator takes to move to the prey, and the ingestion time, there was no significant difference between directly-observed handling times and handling times estimated by fitting functional response curves (Poole et al., 2007). The prey-searching techniques of *M. albidus* and *M. viridis* have previously been described as “[relying] on bumping into … prey during the course of … meanderings” (Fryer, 1957a). This description of rather inefficient prey searching is consistent with our handling time estimates of approximately 30 min to 1 h (Fig. 2c, Table S4).

We found that *M. viridis* was a significantly more efficient predator of *Ae. albopictus* than *M. albidus* (Tables 1 & S5, Fig. 3). An analysis of the gut contents of English Lake District copepods previously found that 15.7% of *M. viridis* had consumed dipterous larvae, compared to only 7.4% of *M. albidus* (Fryer, 1957b). Thus, it is possible that English *M. viridis* may be better adapted to feeding on mosquito larvae than English *M. albidus*. A study using *M. albidus* and *M. viridis* from Northern Ireland against *Culex* mosquitoes from Surrey, UK suggested that *M. albidus* is more effective than *M. viridis* as a biocontrol (Cuthbert et al., 2018b). It appears this is not the case for invasive *Ae. albopictus* prey. A previous study showed that an aedine species (*Ae. aegypti*) was more vulnerable to predation by *M. viridis* than *Cx. pipiens*, despite the larger head capsule width of *Ae. aegypti* (Blaustein and Margalit, 1994). In addition, *M. viridis* was among three cyclopoid copepod species from Northern Ireland that displayed a preference for *Ae. albopictus* larvae over *Cx. pipiens* prey (Cuthbert et al., 2019b). Higher predation efficiency against *Aedes* than against *Culex* may be due to differences in larval anatomy and activity patterns. Previous work has attributed higher survival of *Culex* larvae to lower levels of motility, and to their long bristles, or spines, that either prevent copepod attacks from being successful, or deter attacks entirely by creating the illusion of larger size (Dieng et al., 2003, Marten and Reid, 2007, Soumare and Cilek, 2011). Another study of *M. albidus* and *M. viridis* from Northern Ireland as predators of both *Paramecium caudatum* and *Cx. pipiens* reported that *M. viridis* consumed more prey organisms overall than *M. albidus* (Cuthbert et al., 2019a).

The larger size of *M. viridis* contributes to the higher predation efficiency observed among that species against *Ae. albopictus* (Fig. 3b). Our analysis shows that copepod body mass was a better predictor of predation efficiency than species identity (Tables 1–3). The linear regression model with the highest adjusted R-squared value, 16%, was the model with copepod body mass as the sole predictor of predation efficiency (Tables 3 & S6). This adjusted R-squared value is still low, indicating substantial variation within the predation interaction. The positive relationship we observed between copepod body mass and predation efficiency (Tables 2 & S6, Fig. 3b) is consistent with the findings of a previous meta-analysis on crustacean predation of immature fish, which showed that the predation rate was negatively related to the prey/predator size ratio (Paradis et al., 1996). Our data are also in agreement with the positive linear relationship between body mass and ingestion rate that has been demonstrated in other taxa, including herbivorous and carnivorous endotherms, as well as carnivorous poikilothermic tetrapods (Peters, 1983). Although a recent study using *Ae. japonicus* larvae as prey found that *M. viridis* predation rates began to decrease when predators exceeded 1.8 mm in length (Früh et al., 2019), our data represent copepods of up to 2.5 mm in length, and there is no evidence of lower predation efficiency among the largest predators (Fig. 3b). According to the theory that, at equilibrium, “the rate of food intake is equal to the rate at which food is leaving the gut” (Charnov and Orians, 1973), our results suggest that *M. viridis* may have a higher gut clearance rate than *M. albidus*, possibly due to its larger size.

Despite the higher predation efficiency observed among *M. viridis* (Tables 1 & S5, Fig. 3), we did not detect any differences between the two copepod species in their functional response parameter estimates, when controlling for temperature (Fig. 2b & 2c, Table S4); this was perhaps because the predation efficiency experiment had a larger sample size of observations for each combination of predictors, and therefore more power to detect differences. In addition, while the size difference between *M. albidus* and *M. viridis* was significant in both experimental designs (Supporting Information, Fig. S4), there was a greater difference in size due to species among the copepods used in the predation efficiency experiment than among the copepods used in the functional response experiment.

Although we did not detect any difference in functional response parameter estimates or predation efficiency due to temperature, the temperature at which copepods are cultured impacts their size (Laybourn-Parry et al., 1988), and our results show that copepod size is a determinant of their predation efficiency. A previous study, in which UK *M. albidus* and *M. viridis* were cultured at six different temperatures between 5 and 20°C, showed a consistent negative relationship between adult body mass of both species and culturing temperature (Laybourn-Parry et al., 1988). Evidence of this relationship between size and rearing temperature had already been recorded for freshwater copepods collected in Paris (Coker, 1933). Although cultures tend to have higher rates of reproduction at higher temperatures (Laybourn-Parry et al., 1988), we recommend that the importance of individual body mass be taken into consideration when determining the optimal temperature for mass copepod culturing.

Fecundity data from older studies (Laybourn-Parry et al., 1988, Maier, 1994) have recently been incorporated into metrics used to compare the efficacy of different copepod species as biocontrol agents (Cuthbert et al., 2019b, Cuthbert et al., 2018b). Caution is recommended when interpreting these metrics if the fecundity variable is either based on an inadequate number of observations or derived from multiple populations of copepods. For example, a previous calculation used to determine the “reproductive effort” of *M. viridis* incorporated a clutch weight value based on the dimensions of a single egg sac from a female collected at an unrecorded temperature in South Germany, female body weight data from a population in South Germany, and embryonic development time data from English Lake District copepods held at 20°C (Abdullahi and Laybourn-Parry, 1985, Maier, 1994). This reproductive effort value was then applied to calculate a metric for *M. viridis* from Northern Ireland that were cultured at 25°C (Cuthbert et al., 2019b). While it is time-consuming to collect robust and relevant fecundity data experimentally, biocontrol metrics can only be meaningful if they are based on appropriate original data.

Life history characteristics other than fecundity, particularly lifespan and resistance to starvation, also impact the efficiency of biocontrol agents, and these characteristics have been shown to positively correlate with cyclopoid copepod body size (Dieng et al., 2003, Maier, 1994). The relationship between lifespan and body size is recognized as a general rule in ecology and has been exemplified by comparing the lifespan of an elephant to that of a mouse (Brown et al., 2004). *M. viridis* females from English Lake District cultures have previously been observed to spend approximately 120 days (at 15°C) and 80 days (at 20°C) in the adult life stage (Laybourn-Parry et al., 1988). The fecundity and long-term stability of field copepod populations can be difficult to predict. A year-round field study conducted in Florida, USA of cyclopoid copepod populations in tire piles found that the number of copepods observed was influenced by the age and amount of leaf litter in each tire (Schreiber et al., 1996). In addition, cyclopoid copepod populations survive longer in tires near shade or vegetation because the water in the tire is less likely to evaporate (Marten et al., 1994). In the UK, where *Ae. albopictus* adults are only expected to be active from May to September (Medlock et al., 2006), the stability of year-round copepod populations in tires may not be a priority. The use of copepods as biocontrol agents in UK field sites might benefit from bi-monthly re-applications of well-maintained laboratory cultures, and perhaps more frequent reinforcements in case of long droughts followed by substantial precipitation. A control strategy that is based on precipitation patterns and the estimated survival time of adult copepods in the field would be less risky than a strategy that relies on the viability of subsequent copepod generations.

As the invasion of *Ae. albopictus* into the UK progresses, it may become more relevant to conduct predation experiments on mosquito larvae that are hatched from colonies acclimated to lower temperatures. In our experiments, the *Ae. albopictus* larvae were from a colony that was reared under conditions experienced on warm summer days in Montpellier, France. This approach is currently valid because *Ae. albopictus* has not yet established in the UK and eggs that could be introduced via used tire shipments would be naïve to UK weather conditions. However, in future decades, it may be more appropriate to use *Ae. albopictus* larvae from a colony reared under conditions reflective of the diurnal temperature range experienced in the southern UK.

## Author Contributions

LJC and MCR conceived the ideas and designed the methodology of the predation experiments; AQ, LJC, and MCR collected the data; CGW provided critical advice on cultivating the copepods; AQ improved the *Ae. albopictus* hatching procedure; MCR analyzed the data and led the writing of the manuscript. All authors contributed critically to the drafts and gave final approval for publication.

## Acknowledgements

Many thanks to Flora Dickie, Dr. Maria Hołyńska, and Dr. Ross Cuthbert for their advice and assistance. In addition, we express our gratitude to Dr. Samraat Pawar for his guidance on the functional response literature. This project has received resources funded by the European Union’s Horizon 2020 research and innovation program under grant agreement No 731060 (Infravec2). The work was primarily funded by a President’s PhD Scholarship from Imperial College London awarded to Marie C. Russell.

## Supporting Information

### Materials and methods

#### Collection of field temperature data

In late February and early March of 2018, six empty car tires, each leaning on a wooden crate, were placed in six different locations (Fig. S1), spread between two main sites: South Kensington (London, UK) and Silwood Park (Berkshire, UK). Although most tires were at ground-level near vegetation or shade, the “First Floor Balcony” tire (Fig. S1) had more direct sun exposure, and thus, may be more representative of the conditions present in large tire piles. Each tire was surveyed once a week from the 16^th^ of April until the 30^th^ of October between 11am and 5pm, which matches the time period of predation for both our functional response and predation efficiency experiments. If water was present in a tire, the water temperature was taken using a hand-held Hanna Instruments thermometer (model name: “Checktemp 1”). Temperature data from Silwood Park and South Kensington were recorded from May through September of 2018 (Fig. S2), the months of predicted *Ae. albopictus* adult activity (Medlock et al., 2006). There were 105 observations in total; 16 temperature recordings were missing because the tire was dry, and two were excluded because they were recorded outside the time period of 11am to 5pm.

**Fig. S1.**
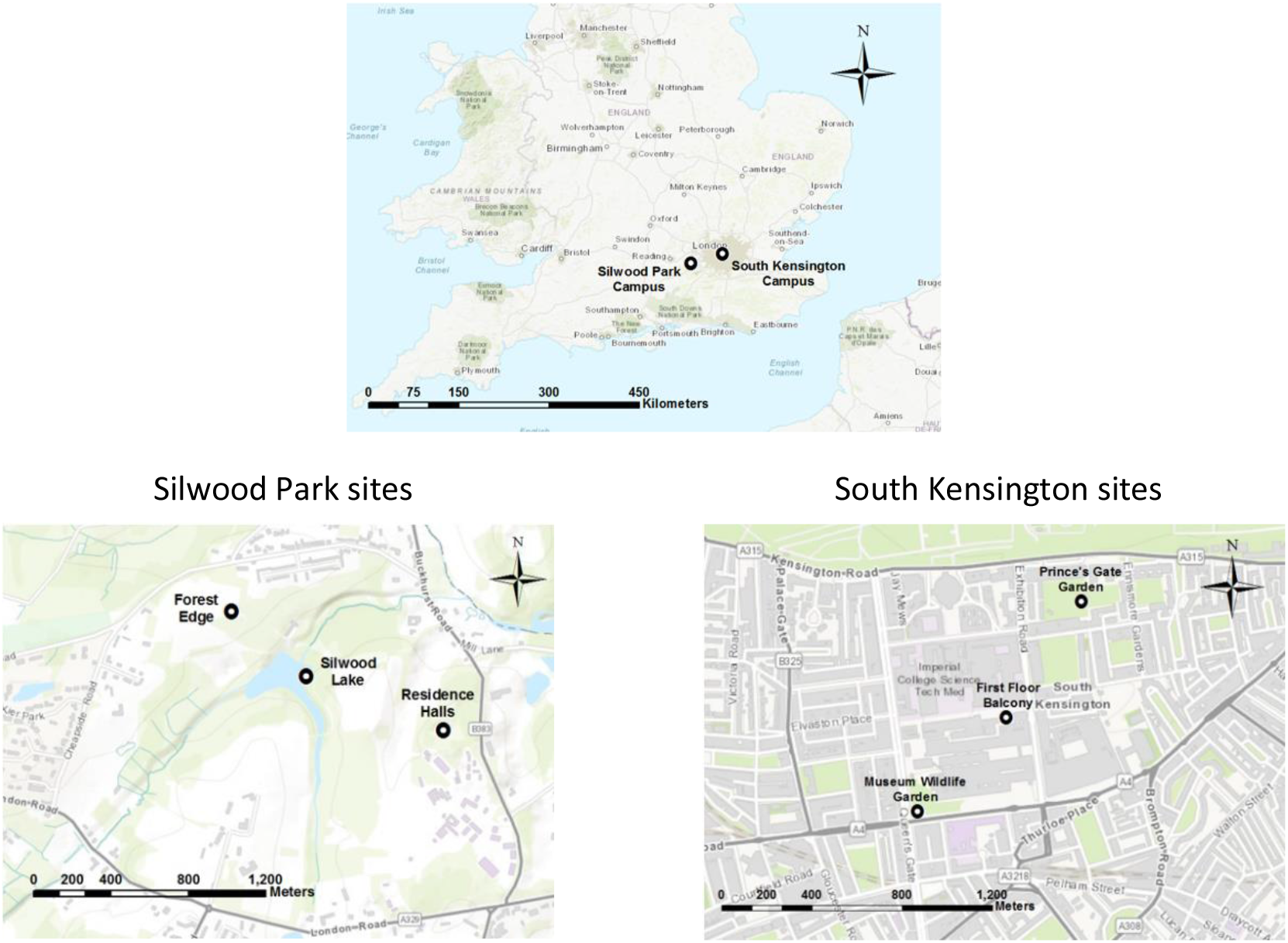
Tire locations

**Fig. S2.**
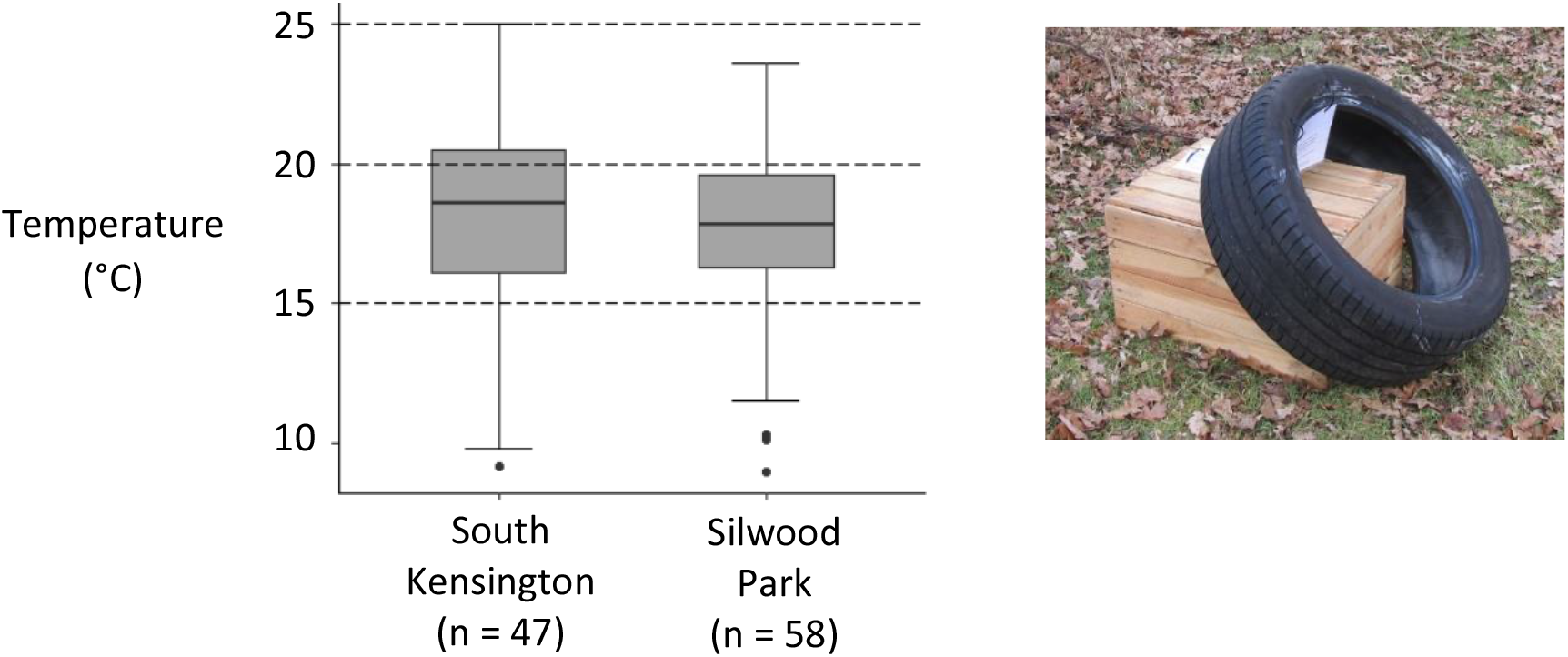
May through September of 2018 weekly tire water temperatures (Dotted lines represent the three temperatures tested in the functional response and predation efficiency experiments.)

**Fig. S3.**
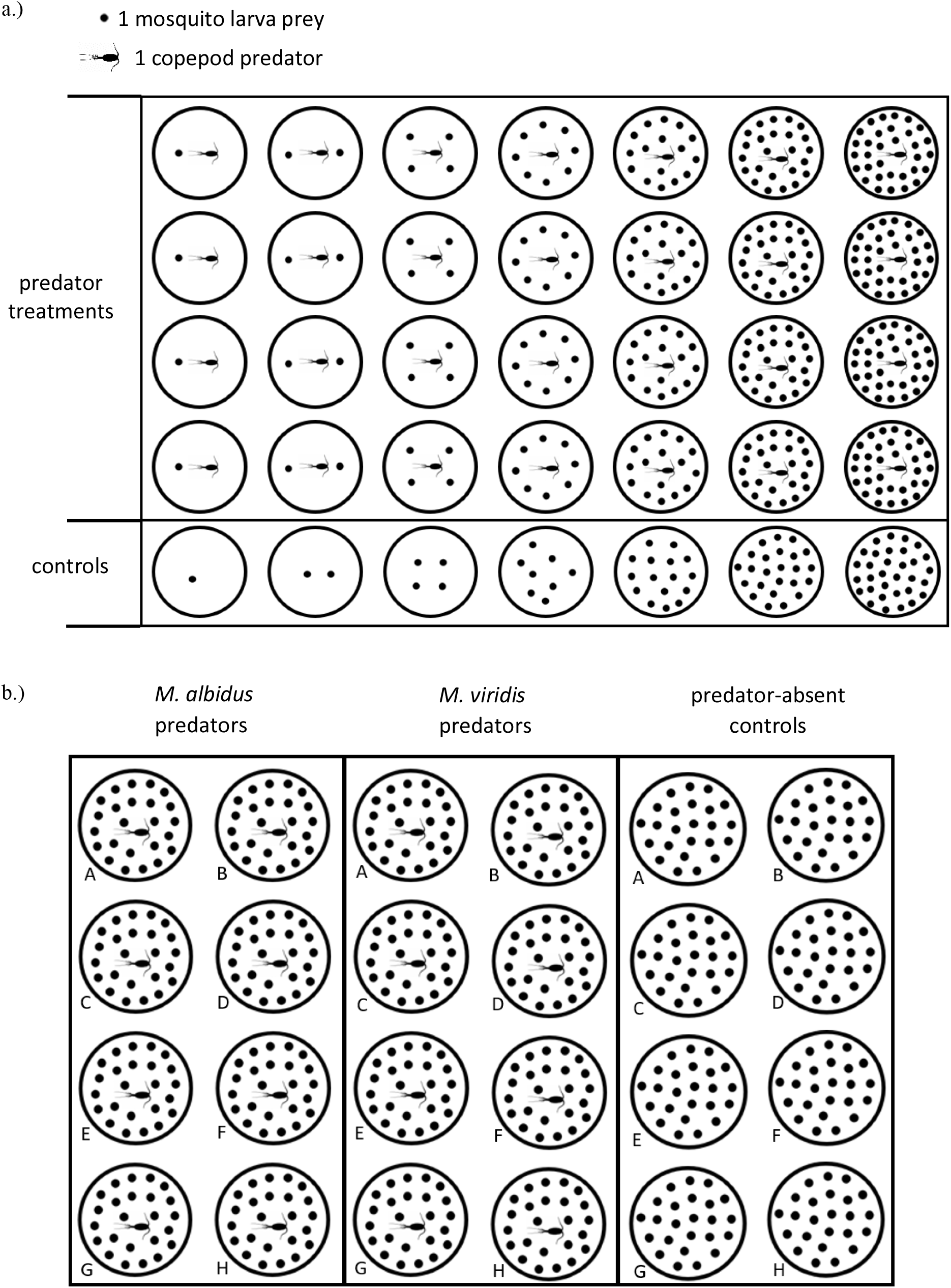
Experimental design: (a) functional response repeated six times, once for each combination of predator species and temperature; (b) predation efficiency repeated three times, once for each temperature (Source of copepod silhouette accessed August 2019: http://phylopic.org/name/f79c5b86-7b73-468a-9ad4-995646398f99)

#### *Ae. albopictus* hatching procedure for functional response and predation efficiency experiments

*Ae. albopictus* eggs were collected from the colony on filter papers and stored in plastic bags containing damp paper towels to maintain humidity. Stored egg papers were submerged in 3 mg/L nutrient broth solution (Sigma-Aldrich © 70122 Nutrient Broth No 1) and oxygen was displaced by vacuum suction for 30 min. Immediately following oxygen displacement, ground fish food (Cichlid Gold Hikari^®^, Japan) was added *ad libitum*. The eggs were left in this solution at 27 ± 1°C for 12 h (Hanson and Craig, 1994) before the newly-hatched larvae were counted (Fig. 1). The hatching temperature was kept high, relative to the experimental temperatures (15-25°C), in order to maximize the hatch rate over a short, semi-synchronous time period, especially since *Ae. albopictus* often hatch in multiple installments (Hanson and Craig, 1994). The larval densities used in the functional response experiments were chosen based on previous studies (Cuthbert et al., 2018, Cuthbert et al., 2019).

#### Predation efficiency linear regression models

The following linear regression model was fitted to test temperature setting and copepod species as predictors of predation efficiency:

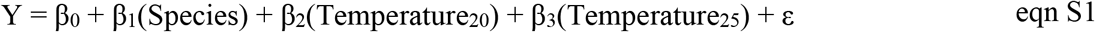

where Y represents the predation efficiency; β_0_ is the Y-intercept; Species is a binary variable denoted as 0 for *M. albidus* and 1 for *M. viridis*; Temperature_20_ is a binary variable denoted as 0 for 15°C and 1 for 20°C; Temperature_25_ is a binary variable denoted as 0 for 15°C and 1 for 25°C; and ε is a random error term, assumed ~N(0, σ^2^). Forty-seven observations were included in the model; one was excluded because the copepod predator (*M. viridis* at 25°C) was found to have died at some point during the 6 h predation period. The statistical significance of each independent parameter estimate was analyzed at α = 0.05.

An additional linear regression model was fitted to test temperature setting and copepod body mass as predictors of predation efficiency:

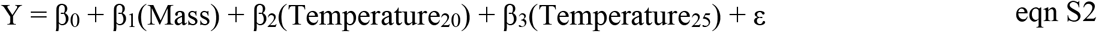

where Mass is a continuous variable referring to each copepod’s body mass in mg; and all other notation is identical to that used in the previous model.

Both linear regression models above were also fitted without any temperature predictors. Akaike information criterion (AIC) values were calculated for model selection, and the Shapiro-Wilk test was used to confirm that the residuals of each model were normally distributed.

### Results

**Table S1.** Results of logistic regression of proportional consumption data as a function of initial prey density using the “frair_test” function

| Species | Temperature (°C) | n<sup>a</sup> | First Order Term<sup>b</sup> | p - value |
| --- | --- | --- | --- | --- |
| *M. albidus* | 15 | 28 | -0.06 | <0.0001 |
| *M. albidus* | 20 | 28 | -0.08 | <0.0001 |
| *M. albidus* | 25 | 25 | -0.04 | 0.0007 |
| *M. viridis* | 15 | 27 | -0.09 | <0.0001 |
| *M. viridis* | 20 | 28 | -0.08 | <0.0001 |
| *M. viridis* | 25 | 25 | -0.09 | <0.0001 |

1.) Controls are not included for determining the type of functional response.
2.) The response is classified as “type II” when there is a significant negative first-order term.

**Table S2.** Results of fitting the “nll.ode.general.mort” function for functional response curves in which background mortality was observed

| Species | Temp.<br>(°C) | n<sup>a</sup> | Parameter | Estimate | Standard<br>Error | p-value | -2 log L |
| --- | --- | --- | --- | --- | --- | --- | --- |
| *M. albidus* | 15 | 35 | attack coefficient | 0.2379 | 0.0921 | 0.0098 | 102.73 |
|  |  |  | handling time | 1.1583 | 0.2782 | <0.0001 |  |
|  |  |  | mortality rate | 0.0063 | 0.0036 | 0.0816 |  |
| *M. albidus* | 20 | 35 | attack coefficient | 0.4561 | 0.1156 | <0.0001 | 90.98 |
|  |  |  | handling time | 0.6036 | 0.0838 | <0.0001 |  |
|  |  |  | mortality rate<sup>2</sup> | 0.0076 | 0.0038 | 0.0426 |  |
| *M. albidus* | 25 | 32 | attack coefficient | 0.4909 | 0.3523 | 0.1634 | 111.25 |
|  |  |  | handling time | 0.9163 | 0.3601 | 0.0109 |  |
|  |  |  | mortality rate<sup>b</sup> | 0.0289 | 0.0100 | 0.0040 |  |
| *M. viridis* | 15 | 34 | attack coefficient | 0.4911 | 0.1214 | <0.0001 | 100.14 |
|  |  |  | handling time | 0.5163 | 0.0705 | <0.0001 |  |
|  |  |  | mortality rate | 0.0038 | 0.0027 | 0.1569 |  |
| *M. viridis* | 20 | 35 | attack coefficient | 0.4511 | 0.1063 | <0.0001 | 93.30 |
|  |  |  | handling time | 0.4848 | 0.0667 | <0.0001 |  |
|  |  |  | mortality rate | 0.0057 | 0.0033 | 0.0823 |  |

a.) Number of empirical observations includes both predator and control treatments.
b.) Background prey mortality had a significant impact on fitting the model for *M.*
*albidus* at 20 and 25°C.

**Table S3.** Results of fitting the “nll.bolker” function for functional response curves in which background mortality was either not observed or insignificant

| Species | Temp.<br>(°C) | n | Parameter | Estimate | Standard<br>Error | p-value | -2 log L |
| --- | --- | --- | --- | --- | --- | --- | --- |
| *M. albidus* | 15 | 28 | attack coefficient | 0.2166 | 0.0716 | 0.0025 | 89.71 |
|  |  |  | handling time | 0.9189 | 0.1696 | <0.0001 |  |
| *M. viridis* | 15 | 27 | attack coefficient | 0.4803 | 0.1137 | <0.0001 | 92.14 |
|  |  |  | handling time | 0.4853 | 0.0619 | <0.0001 |  |
| *M. viridis* | 20 | 28 | attack coefficient | 0.4430 | 0.0989 | <0.0001 | 83.57 |
|  |  |  | handling time | 0.4447 | 0.0554 | <0.0001 |  |
| *M. viridis* | 25 | 25 | attack coefficient | 0.5629 | 0.1438 | <0.0001 | 85.53 |
|  |  |  | handling time | 0.6527 | 0.0725 | <0.0001 |  |

**Table S4.** Functional response parameter estimates and 95% confidence intervals

| Species | Temp. (°C) | Attack Coefficient<br>(attacks per hour) | Handling Time<br>(hours) |
| --- | --- | --- | --- |
| *M. albidus* | 15 | 0.217, (0.116 – 0.425) | 0.919, (0.596 – 1.28) |
| *M. albidus* | 20 | 0.456, (0.277 – 0.747) | 0.604, (0.452 – 0.798) |
| *M. albidus* | 25 | 0.491, (0.178 – 13.9) | 0.916, (0.451 – 1.99) |
| *M. viridis* | 15 | 0.480, (0.305 – 0.777) | 0.485, (0.368 – 0.615) |
| *M. viridis* | 20 | 0.443, (0.287 – 0.691) | 0.445, (0.337 – 0.559) |
| *M. viridis* | 25 | 0.563, (0.338 – 0.921) | 0.653, (0.519 – 0.808) |

**Table S5.** Linear regression of predation efficiency by copepod species (n = 47)

| Parameter | Estimate | Standard Error | p-value | Adjusted R<sup>2</sup> | AIC |
| --- | --- | --- | --- | --- | --- |
| Intercept | 19.97 | 2.314 | $4.25 \times 10^{-11}$ | 0.081 | 362.6 |
| Species (*M. viridis*, ref = *M. albidus*) | 7.42 | 3.308 | 0.0299 |  |  |

**Table S6.** Linear regression of predation efficiency by copepod body mass (n = 47)

| Parameter | Estimate | Standard Error | p-value | Adjusted R<sup>2</sup> | AIC |
| --- | --- | --- | --- | --- | --- |
| Intercept | 11.19 | 4.33 | 0.0130 | 0.156 | 358.5 |
| Body Mass (mg) | 34.75 | 11.26 | 0.0035 |  |  |

#### Copepod body sizes by experimental design and species

The ten gravid female copepods of each species that were measured as a validation set for the size of adult females included in the functional response experiments ranged from 1.2 to 1.7 mm for *M. albidus*, and from 1.4 to 2.3 mm for *M. viridis*. The non-gravid copepods used as predators in these experiments ranged from 1.2 to 1.9 mm (n = 81) for *M. albidus*, and from 1.4 to 2.5 mm (n = 80) for *M. viridis*. The eighty-one *M. albidus* copepods included in the functional response curve experiments ranged from 0.09 to 0.32 mg in body mass (median = 0.20 mg), and the eighty *M. viridis* copepods included in the functional response curve experiments ranged from 0.14 to 0.67 mg in body mass (median = 0.32 mg). Results of the Shapiro-Wilk test showed that the distribution of body masses included in the functional response experiments was significantly different from a normal distribution for both *M. albidus* (p-value = 0.0019) and *M. viridis* (p-value = 0.0001). The results of a Wilcoxon rank sum test showed that *M. viridis* used in the functional response experiments were generally larger than *M. albidus* used in the same experiments (p-value < 0.0001, Fig. S4).

Twenty-four *M. albidus* copepods were used as predators in the predation efficiency experiments, ranging in body mass from 0.14 to 0.36 mg (mean = 0.24 mg, standard deviation = 0.06 mg), and 23 *M. viridis* were used, ranging in body mass from 0.23 to 0.67 mg (mean = 0.48 mg, standard deviation = 0.10 mg). Results of the Shapiro-Wilk test showed that the distribution of body masses included in the predation efficiency experiments was not significantly different from a normal distribution for both *M. albidus* (p-value = 0.0665) and *M. viridis* (p-value = 0.2128). The results of a Welch two sample t-test for samples of unequal variance showed that *M. viridis* used in the predation efficiency experiments were significantly larger than *M. albidus* used in the same experiments (p-value < 0.0001, Fig. S4). The Wilcoxon rank sum test was used to identify differences in copepod body mass by experimental design, controlling for species. Copepods used in the predation efficiency experiment were generally larger than those used in the functional response experiment for both *M. albidus* (p-value = 0.0008) and *M. viridis* (p-value = 0.0003).

**Fig. S4.**
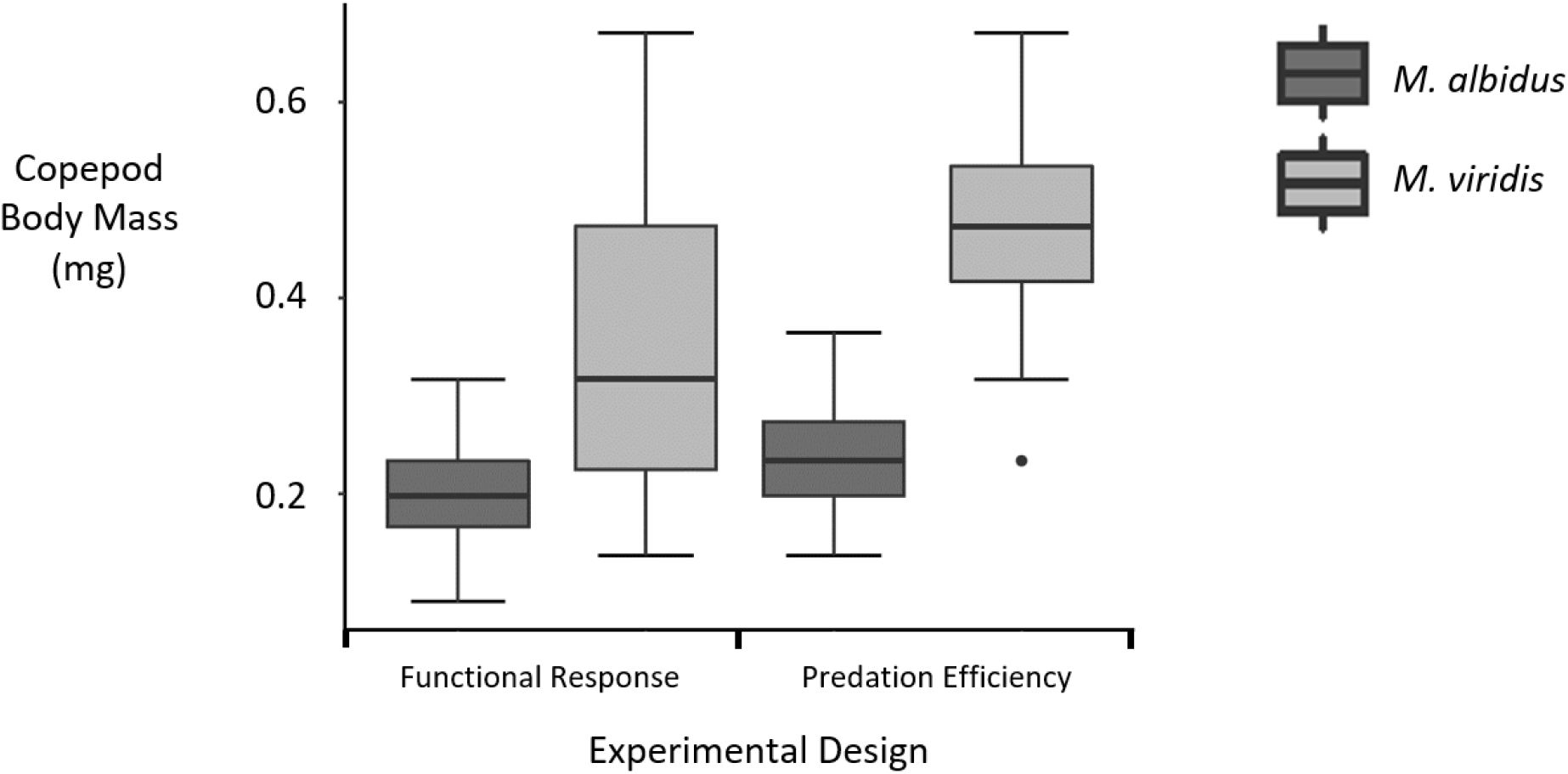
Boxplot of copepod body mass by experimental design. Sample sizes: Functional response, *M. albidus* = 81; Functional response, *M. viridis* = 80; Predation efficiency, *M. albidus* = 24; Predation efficiency, *M. viridis* = 23.

